# Transient exposure to oxygen or nitrate reveals ecophysiology of fermentative and sulfate-reducing benthic microbial populations

**DOI:** 10.1101/146886

**Authors:** Zainab Abdulrahman Beiruti, Srijak Bhatnagar, Halina E. Tegetmeyer, Jeanine S. Geelhoed, Marc Strous, S. Emil Ruff

## Abstract

**Conflict of interest:** The authors declare no conflict of interest.

**Short summary:** Fermentation coupled to sulfate reduction is a globally important process for the remineralization of organic carbon in marine sediments. The present study uses long-term, replicated continuous culture bioreactors and meta-omics to investigate the ecophysiology of the involved microbial populations at an unprecedented resolution. We reveal complex trophic networks, in which fermenters and sulfate reducers coexist with nitrate- and oxygen respirers, we indicate strategies and niches of the microbial populations, and describe a novel and widespread, yet uncultured fermentative organism. These insights are crucial to understand fermentation coupled to sulfate reduction and relevant to assess microbial dynamics and community-level responses in coastal ecosystems.

For the anaerobic remineralization of organic matter in marine sediments, sulfate reduction coupled to fermentation plays a key role. Here, we enriched sulfate-reducing/fermentative communities from intertidal sediments under defined conditions in continuous culture. We transiently exposed the cultures to oxygen or nitrate twice daily and investigated the community response. Chemical measurements, provisional genomes and transcriptomic profiles revealed trophic networks of microbial populations. Sulfate reducers coexisted with facultative nitrate reducers or aerobes enabling the community to adjust to nitrate or oxygen pulses. Exposure to oxygen and nitrate impacted the community structure, but did not suppress fermentation or sulfate reduction as community functions, highlighting their stability under dynamic conditions. The most abundant sulfate reducer in all cultures, related to *Desulfotignum balticum*, appeared to have coupled acetate oxidation to sulfate reduction. We described a novel representative of the widespread uncultured phylum *Candidatus* Fermentibacteria (formerly candidate division Hyd24-12). For this strictly anaerobic, obligate fermentative bacterium, we propose the name *Ca.* “Sabulitectum silens” and identify it as a partner of sulfate reducers in marine sediments. Overall, we provide insights into the metabolic network of fermentative and sulfate-reducing microbial populations, their niches, and adaptations to a dynamic environment.

## Introduction

Around 30% of the total oceanic phytoplankton-derived primary production occurs along the continental margins (Walsh, 1991) and up to 50% of this organic matter reaches the surface of shallow coastal sediments. This organic matter can be re-mineralized by the microorganisms in the surface sediment using a broad suite of electron acceptors, such as oxygen, nitrate, metal oxides and sulfate (Henrichs and Reeburgh, 1987; Canfield *et al.*, 1993; Janssen *et al*., 2005). It has been estimated that about 50% of the total organic carbon mineralization in shallow sediments (Jørgensen, 1982) and salt marsh sediments (Howes *et al*., 1984) and up to 35% of the total mineralization in intertidal flats (Billerbeck *et al*., 2006) is coupled to sulfate reduction. Yet, despite the global importance of sulfate reduction, the ecophysiology of the involved microorganisms and their environmental controls are poorly constrained.

The sulfate-reducing microbial populations in the surface sediments of intertidal flats are exposed to pulses of oxygen approximately twice daily, because of tidal cycling. In addition, the communities may be regularly exposed to pulses of nitrogen from riverine sources (van Beusekom, 2005; Boyer *et al*., 2006). It is thus very likely that sulfate reducers and also other key anaerobic functional types, such as fermenters, are adapted to these ecosystem dynamics and survive exposure to oxygen and nitrate. Generally, the availability of oxygen leads to a lower relative importance of sulfate reduction, because electron acceptors tend to be consumed in a thermodynamically determined order (the redox cascade). According to this order, oxygen is used first, followed by nitrate, manganese and iron oxides, and finally sulfate (Froelich *et al*., 1979). Hence, sulfate is thought to be the predominant electron acceptor only in the anoxic layers after other electron acceptors are depleted. Sulfate-reducing bacteria are often strict anaerobes and couple the oxidation of molecular hydrogen or organic compounds to the complete reduction of sulfate to sulfide (Muyzer and Stams, 2008; Rabus *et al*., 2013). Nevertheless, sulfate-reducing bacteria were detected throughout the whole sediment of an intertidal flat, including the aerobic and denitrifying zones (Llobet-Brossa *et al*., 2002; Mußmann *et al*., 2005; Gittel *et al*., 2008). In addition, it was found that intertidal flats are a sink for riverine and atmospheric nitrogen (Gao *et al*., 2012), with the microbial conversion of nitrate to ammonium or dinitrogen (Marchant *et al*., 2014) and the internal storage of nitrate in benthic diatoms (Stief *et al*., 2013) being widespread and important processes. Also, nitrite is common in intertidal flats and it was found that some sulfate reducers, like *Desulfovibrio desulfuricans*, are able to grow on hydrogen coupled to ammonification of nitrate or nitrite (Dalsgaard and Bak, 1994). Although, much progress has been made in understanding the key processes and populations in intertidal sediments, e.g. elucidating the environmental controls of nitrate respiration (Kraft *et al*., 2014) and the impact of chemical gradients on community structure (Chen *et al*., 2017), the trophic network defining combined fermentation and sulfate reduction remains largely unknown.

A major challenge in microbial ecology in general is to understand the dynamics of an ecosystem and its impact on the microbial communities (Widder *et al*., 2016). This can be addressed e.g. by investigating the resistance and resilience of microbial communities to perturbations (Shade *et al*., 2012; Lee *et al*., 2017), or by investigating the response of microbial communities to recurring events (Ward *et al*., 2017). Simple model systems are a promising tool to disentangle community dynamics and constrain cause and effect (Widder *et al*., 2016). To investigate the effect of the tidal cycle on fermentation coupled to sulfate reduction as a community function, we set up defined continuous cultures and created a homogeneous microbial habitat that selected for communities of sulfate-reducing and fermentative bacteria. We inoculated the cultures with biomass from tidal flat sediments that were exposed to a tidal cycle. The effect of diurnal exposure to oxygen or nitrate on the microbial activity and community structure was examined by combined chemical, metagenomic and transcriptomic analyses. Using this setup, we gained insights into fermentation coupled to sulfate reduction and the involved trophic networks, as well as into the ecophysiology of an uncultured candidate phylum.

## Results and Discussion

### Physiology of the continuous cultures

Six cultures were inoculated with cell suspensions obtained from intertidal sediments of the Janssand tidal flat. The cultures were continuously supplied with sulfate as electron acceptor and a mixture of glucose, seven different amino acids and acetate as electron donors. This mixture was chosen to stimulate the growth of a wide range of organisms, and represents compounds of decaying biomass, which is the main organic carbon source in marine sediments. After two days, sulfide was detected in all cultures and increased during the first 150 days, to a concentration of 2 – 6 mM (Fig. 1). All six cultures were incubated for 20 days under identical sulfate-only conditions to establish anaerobic communities that carry out fermentation-coupled sulfate reduction. From day 21 onward, four of the cultures were treated with oxygen or nitrate pulses, while two cultures remained untreated. Oxygen was supplied to two replicate cultures (Oxy-1 and Oxy-2) for 30 minutes twice daily, by sparging the cultures with air. Two replicate cultures (Nit-1 and Nit-2) were supplied with a nitrate solution for seven minutes twice daily. The final two replicate cultures (Con-1 and Con-2) did not receive any additional electron acceptor and served as an untreated control. The biomass in each of the cultures remained stable during the entire experiment (OD_600_: ~0.15; Fig. S2). The sulfide concentrations in the cultures, in combination with the nature of the provided carbon sources, indicated that we selected for a syntrophic community of fermenters and sulfate reducers. It was expected that the fermenting bacteria convert glucose and amino acids to short-chain fatty acids, lactate, alcohols, or hydrogen. These could then be used as carbon sources and/or electron donors by the sulfate-reducing bacteria (Rabus *et al*., 2013).

**Figure 1:**
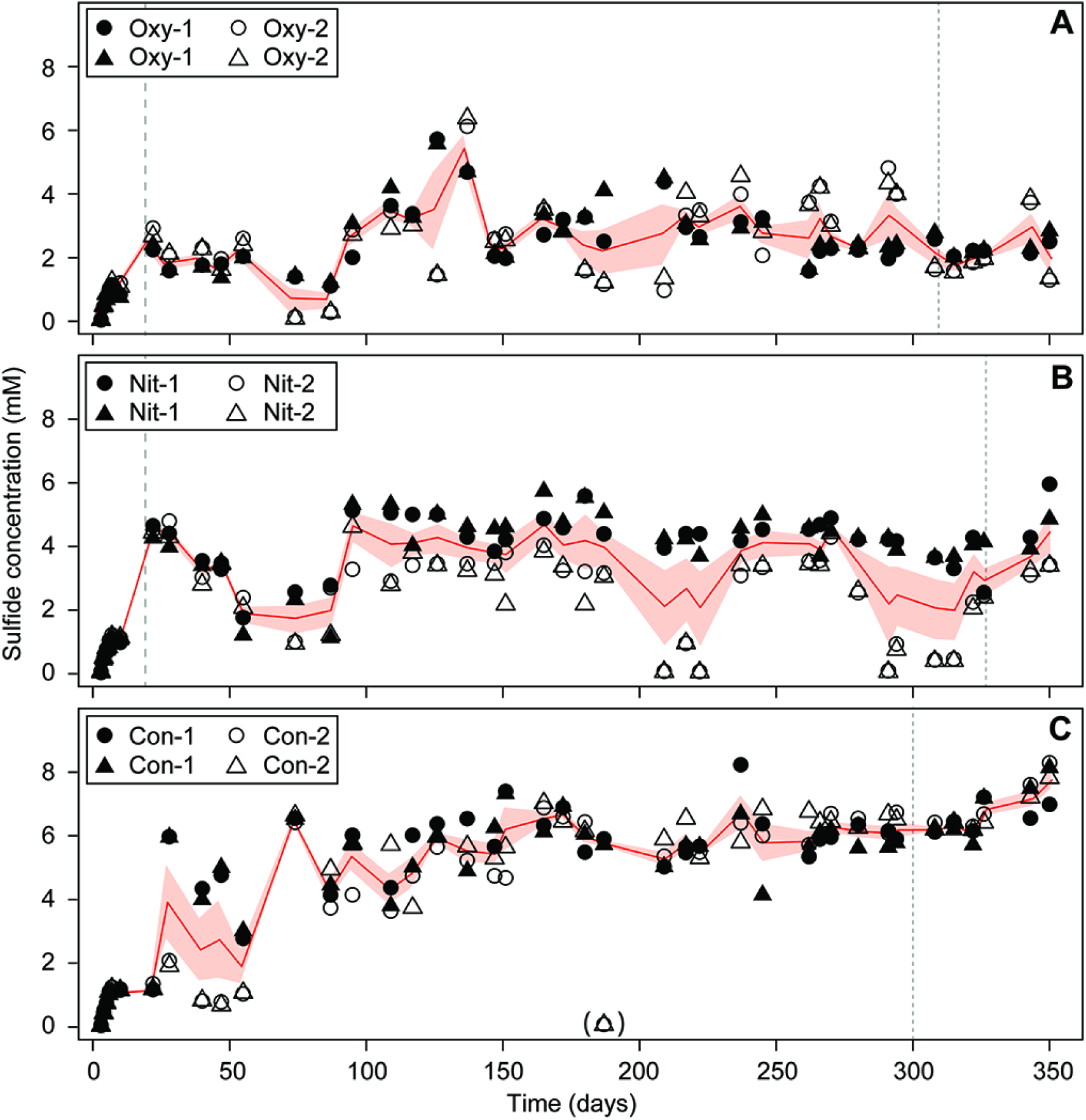
Sulfide concentrations in the replicate cultures treated with oxygen (A), treated with nitrate (B), and in the untreated control with sulfate as sole electron acceptor (C). Duplicate measurements of each culture are shown as triangles and circles, the red line depicts the mean of four measurements, the red ribbon represents standard error of the mean. The start of the treatments is indicated by dashed lines, sampling time points for metagenomics and metatranscriptomics are indicated by dotted lines.

We characterized the cultures in detail on day 311 (Oxy-1 and Oxy-2), day 327 (Nit-1 and Nit-2) and day 300 (Con-1 and Con-2). During the air supply, the oxygen concentration was stable at around 1.3% air saturation (3.1 µM), while the sulfide concentration decreased by 0.7±0.4 mM (Fig. S3A). In the cultures supplied with nitrate, sulfide concentrations did not decrease (Fig. S3B) and the nitrate was metabolized within ~200 min after termination of the supply (Fig. S3D). In both treatments we observed the transient production of elemental sulfur. In the oxygen treatment we measured sulfur concentrations of up to 0.8 mM immediately after the start of aeration, decreasing to ~0.1 mM within 2–4 hours (Fig. S3E). In the nitrate cultures, sulfur was increasing from ~0.1 mM to up to 0.4 mM within 2 hours, and decreased to ~0.1 mM within 4 hours after the start of the treatment (Fig. S3 F). Using ^15^ N-labeled nitrate we found no production of ^15^ N-labeled N2, which indicated that ammonia may have been the end product of nitrate reduction. Ammonia production could not be assessed directly because of the high background ammonia concentration that resulted from ammonification of the supplied amino acids. Over the one year incubation, transient oxygen supply yielded the lowest average sulfide concentrations (2.3±0.3 mM; Fig. 1A), followed by the cultures that received nitrate (4.2±0.6 mM, Fig. 1B) and the untreated control cultures (6.3±0.7 mM, Fig. 1C). Fluctuations in sulfide concentration were highest in the nitrate treatment and lowest in the cultures that were not exposed to oxygen or nitrate. Yet, the cyclic exposure to oxygen or nitrate did not suppress sulfide production (Fig. S3), and thus sulfate reduction, as a community function. Aerobic respiration and ammonification coincided with a decreased magnitude and stability of the sulfide concentration, likely due to microbial re-oxidation of sulfide, or due to competition between sulfate reducers, aerobes or nitrate ammonifiers.

### Microbial communities and their response to cyclic exposure to oxygen and nitrate

After around 300 days of cultivation, we sequenced the metagenomes of the six continuous cultures. We hypothesized that cyclical exposure to oxygen or nitrate alters resource access to create ecological niches that resemble those present in permeable intertidal sediments. Each treatment would thus select for a different microbial community. Indeed, the community structure was different between the treatments (Fig. 2, S4). Yet, all treatments and cultures had a similar microbial community composition (Fig. 2). The nitrate treatment favored fermentative organisms that were less abundant in other treatments, such as *Defluviitaleaceae* (bin K) and certain *Spirochaeta* (bin M/bin N) (Fig. 2, 4A, S5). Moreover, each of the replicate cultures Nit-1 and Nit-2 selected for communities of different structure, although they experienced the same selective pressure. The cultures exposed to oxygen were also different from each other, based on the relative abundances of *Clostridia* (bin H, bin J) and *Psychromonas* (bin B). In contrast, the communities in the untreated replicate controls Con-1 and Con-2 had a nearly identical community structure after 300 days of cultivation (Fig. 2, S4). Despite the different communities in the nitrate and oxygen treated cultures, fermentation coupled to sulfate reduction was not greatly affected as a community function, as inferred from gene expression (Fig. 3, Dataset 1) and the production of sulfide (Fig. 1). This functional similarity may be explained by the presence of fermentative populations that are phylogenetically different, but perform similar metabolisms (Allison and Martiny, 2008).(Burke *et al*., 2011)

**Figure 2:**
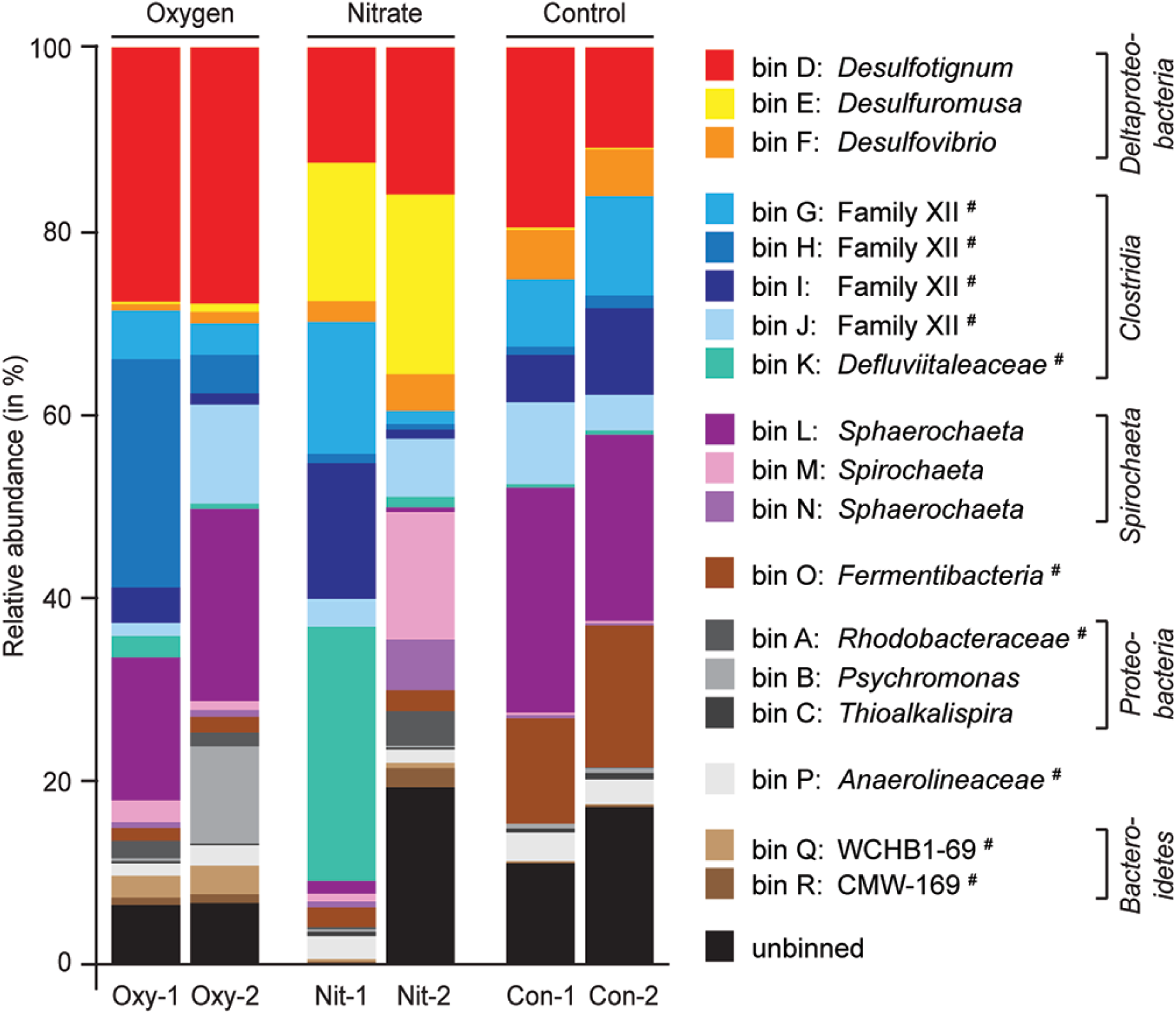
Estimated relative abundances of bins in cultures treated with oxygen (Oxy-1, Oxy-2), with nitrate (Nit-1, Nit-2) and in an untreated control (Con-1, Con-2). The bins were classified to genus level. Populations that affiliated with genera lacking a cultured representative are marked with #. For these bins we reported the closest taxonomically assigned, phylogenetic level, e.g. bin K affiliates with an uncultured genus in the family *Defluviitaleaceae*. Taxonomic assignment is based on the SILVA small ribosomal subunit reference database (SSURef, v123). Relative abundances were obtained by mapping metagenomic sequence reads to the assembled contigs of each bin. The phylogeny of most bins is provided (bin O: Fig. 6, bin A-N: Fig. S6-S9).

**Figure 3:**
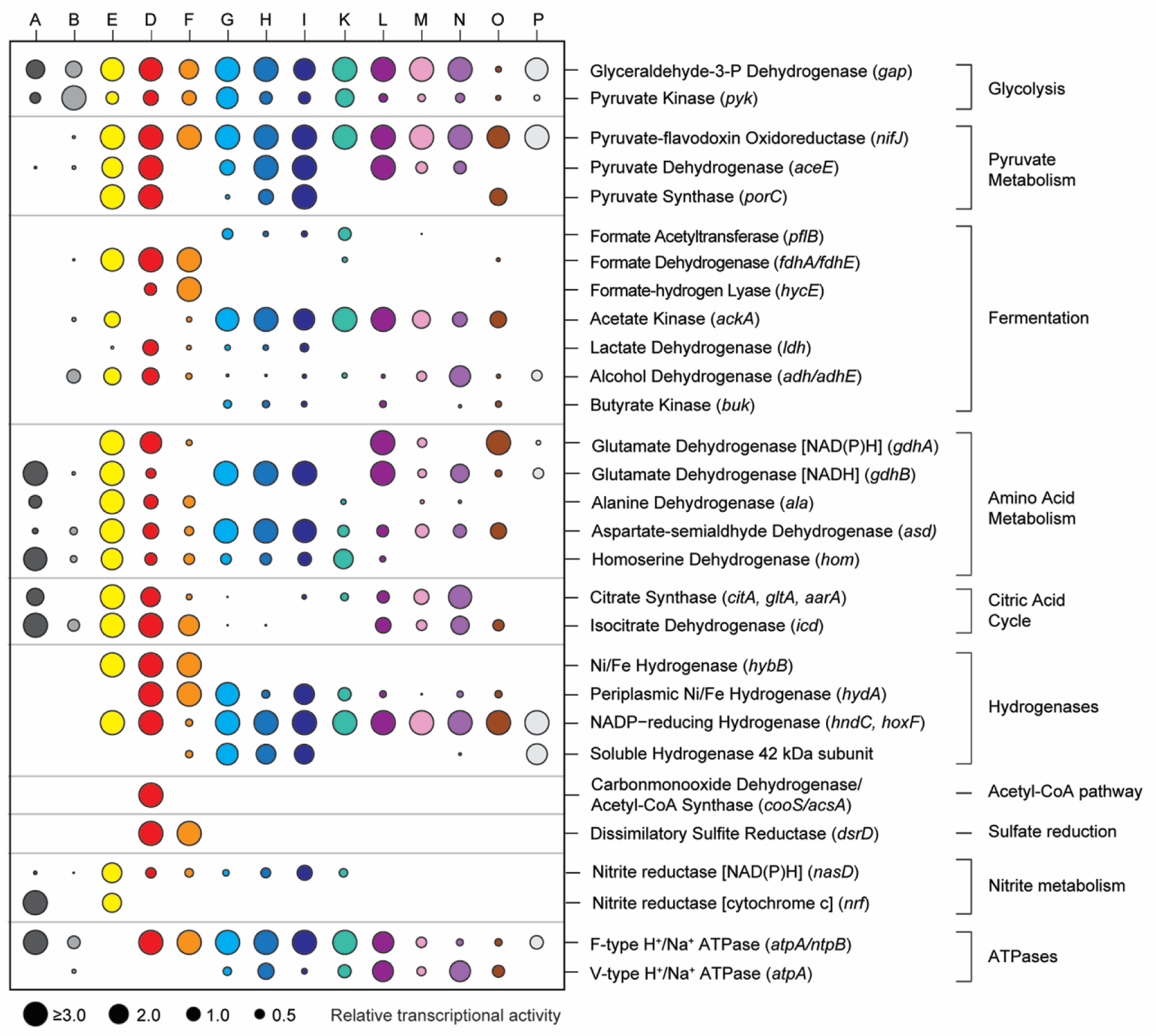
Metabolic capabilities of the populations based on key genes. Circle size represents relative transcriptional activity averaged over all ten transcriptomes. The absence of a circle shows that gene transcription was not detected in any of the treatments and cultures. Note: For reasons of visualization all relative transcriptional activities above 3.0 are shown as ≥3.0.

To study the communities in detail, we focused on metagenomic bins with relatively long contigs with relatively equal coverage distribution and a consistent taxonomic signature (Table 1). These bins can be considered provisional genome sequences that represented the genetic repertoire of the most abundant populations (Table 1, Fig. 2), which were all present in the inoculum as well (Table 1). The 16S rRNA gene sequences of fifteen of these bins were used for a detailed analysis of their taxonomic and phylogenetic affiliation (Fig. S6-S9). *Proteobacteria, Firmicutes*, and *Spirochaeta* were the phyla with the overall highest relative abundance (Fig. 2) and dominated the metagenomes of each treatment. Of the 18 most sequence abundant bins, only bin D affiliating with sulfate-reducing *Desulfotignum* had a high relative abundance across all treatments and cultures, while the other 17 bins were either always relatively low abundant or thrived only under certain conditions (Fig. 2). Most organisms were predicted to have a fermentative (bin G-P) or sulfate-reducing (bin D and F) lifestyle (Fig. 3). Bin A-C, E, Q and R belonged to heterotrophs that were selected in the oxygen or nitrate treatments and hence a respiratory lifestyle is most likely. For the bins C (*Thioalkalispira*), Q, and R (*Bacteroidetes*) meaningful metabolic inferences were impossible because the binned metagenomic data was too scarce (Table 1), bin J was not further analyzed due to a high percentage of contamination (40%).

**Table 1.** Properties of the 18 bins obtained from metagenomes of the six continuous cultures (Oxy-1/2, Nit-1/2, Con-1/2)

| Bin | A | B | C | D | E | F | G | H | I |
| --- | --- | --- | --- | --- | --- | --- | --- | --- | --- |
| Affiliation | Rhodo<br>bacterales | Altero<br>monadales | Chromati<br>ales | Desulfo<br>bacterales | Desulfuro<br>monadales | Desulfo<br>vibrionales | Firmi<br>cutes | Clostridi<br>ales | Clostridi<br>ales |
| Size (Mb) | 3.85 | 3.08 | 1.55 | 4.77 | 4.10 | 4.11 | 4.27 | 4.45 | 4.59 |
| Number of contigs | 3272 | 1897 | 1650 | 594 | 237 | 885 | 2310 | 351 | 562 |
| N50 contig length (kb) | 1.5 | 2.5 | 1.1 | 126 | 58.9 | 14.6 | 2.8 | 27.6 | 107 |
| GC content (%) | 60.1 | 50.6 | 47.6 | 51.9 | 50.2 | 53.9 | 40.7 | 37.6 | 39.9 |
| Number of CSCGs | 131 | 115 | 71 | 132 | 131 | 138 | 121 | 112 | 119 |
| Number of tRNAs | 41 | 34 | 16 | 43 | 50 | 60 | 62 | 31 | 51 |
| Completeness (%) | 71.1 | 73.2 | 33.5 | 84.9 | 89.3 | 86 | 72.9 | 74.2 | 79.4 |
| Contamination (%) | 25 | 7 | 3 | 15 | 11 | 8 | 9 | 8 | 20 |
| *In situ* relative abundance,<br>Mean ± S.D. (%) | 0.05±0.026 | 0.035±0.022 | 0.013±0.009 | 0.041±0.024 | 0.048±0.028 | 0.062±0.036 | 0.002±0.001 | 0.02±0.012 | 0.031±0.02 |

| Bin | J | K | L | M | N | O | P | Q | R |
| --- | --- | --- | --- | --- | --- | --- | --- | --- | --- |
| Affiliation | Clostridi<br>ales | Clostridi<br>ales | Spiro<br>chaetales | Spiro<br>chaetales | Spiro<br>chaetales | Fermenti<br>bacteria | Anaero<br>linea | Bacteroi<br>detes | Bacteroi<br>detes |
| Size (Mb) | 6.69 | 3.86 | 3.36 | 3.57 | 3.78 | 2.92 | 2.51 | 1.68 | 2.30 |
| Number of contigs | 3459 | 248 | 40 | 317 | 691 | 96 | 1147 | 1661 | 2657 |
| N50 contig length (kb) | 3.3 | 118 | 144 | 160 | 10.1 | 222 | 6.0 | 1.2 | 0.99 |
| GC content (%) | 34.5 | 45.4 | 53.5 | 36.8 | 35.7 | 56.6 | 52.8 | 43.5 | 42.1 |
| Number of CSCGs | 182 | 126 | 129 | 88 | 93 | 98 | 128 | 69 | 100 |
| Number of tRNAs | 41 | 39 | 47 | 47 | 36 | 39 | 36 | 11 | 11 |
| Completeness (%) | 86.1 | 90.7 | 88 | 82.1 | 75.3 | 76.9 | 86.7 | 40.5 | 42.4 |
| Contamination (%) | 40 | 18 | 10 | 15 | 17 | 5 | 9 | 2 | 21 |
| *In situ* relative abundance,<br>Mean ± S.D. (%) | 0.019±0.01 | 0.023±0.011 | 0.021±0.015 | 0.021±0.015 | 0.016±0.011 | 0.019±0.01 | 0.015±0.009 | 0.005±0.004 | 0.012±0.008 |

CSCG: Conserved single-copy gene S.D.: Standard deviation of the mean

To infer the metabolic activity of key organisms in the cultures and in response to the applied treatment, we sequenced ten metatranscriptomes after 300 (control), 311 (oxygen treatment) and 327 (nitrate treatment) days of incubation. We sampled one hour before the treatment and directly after the treatment subsided (Fig. S3 C, D). This enabled us to sketch a trophic network in the cultures (Fig. 4A) and look at differences in their gene expression caused by the treatments. The transcriptional activity mirrored relative abundance, such that populations that were abundant in a treatment, were also most active. The differences in overall gene transcription before and after the treatment were not very pronounced. The relative transcription of most genes involved in anabolism, catabolism and energy metabolism showed minor changes, suggesting that after 300 days the key organisms were very well adapted to the provided cyclic environment. Consistent and large differences caused by the treatment were mainly detected for genes involved in oxidative (Fig 4B) and general (Fig 4C) stress protection. The genes that were transcribed by the populations indicated that each population had a slightly different strategy to cope with stress.

**Figure 4:**
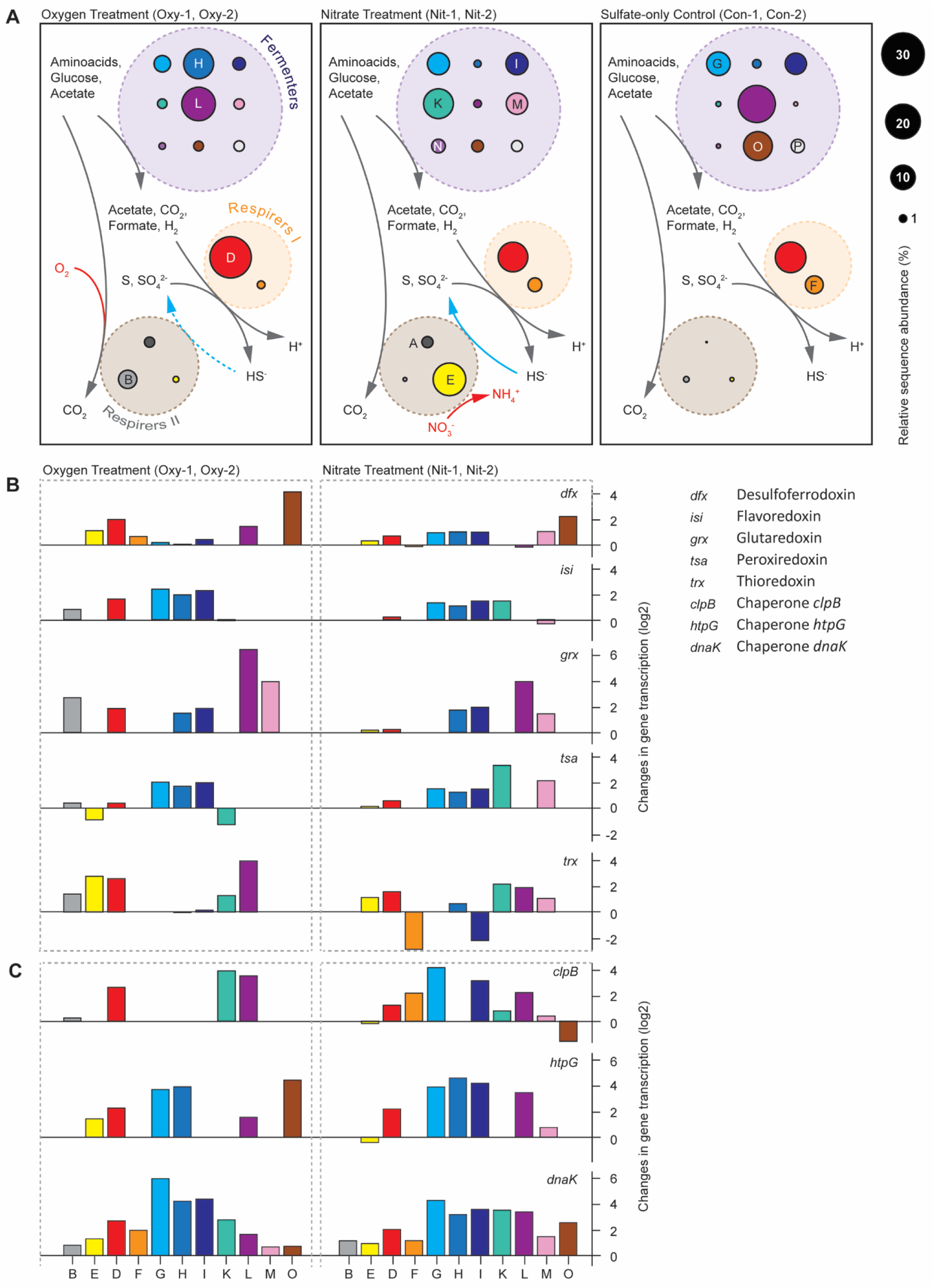
Schematic of the trophic network of key populations, and transcriptional changes of stress response genes. (A) Most abundant obligate fermentative heterotrophs (Fermenters), sulfate-reducing bacteria (Respirers I) and associated respiratory heterotrophs (Respirers II) in the three different conditions. The network is based on metagenomic and /–transcriptomic data. All 14 shown bins were present in all cultures. Circle size represents estimated relative abundance. Only one organism (bin D) was abundant in all cultures. Arrows depict key pathways that occur in all (grey), two (blue) or one condition (red). (B) Change of gene transcription caused by the treatment with oxygen or nitrate. Values are log2-transformed ratios of gene transcription in replicate cultures after and before the treatment, i.e. a value of 1 means that gene transcription was twice as high after the treatment than before the treatment; a value of -2 means a four-fold decrease in transcription.

#### Sulfate reducers

Most of the sulfate reduction was likely performed by *Deltaproteobacteria* affiliating with *Desulfotignum balticum* (bin D, Fig. S6) and *Desulfovibrio profundus* (bin F, Fig. S6). Both organisms constitutively expressed bd-type terminal oxidases to respire oxygen and protect oxygen-sensitive enzymes (Ramel *et al*., 2013). Genes encoding for sulfate adenylyltransferase (*sat*), adenylyl-sulfate reductase (*aps*) and dissimilatory sulfite reductase (*dsr*) were also transcribed by both organisms (Fig. 3, Dataset 1). *Desulfotignum* dominated all conditions based on relative abundance, yet *Desulfovibrio* seemed to have a higher relative transcription of *dsr* genes than *Desulfotignum* in the nitrate-supplied and untreated cultures (Fig. S5). *Desulfotignum* transcribed NiFe(Se)-hydrogenases (e.g. *hyb*) and c-type cytochromes, which are needed to use hydrogen as an electron donor (Heidelberg *et al*., 2004). *Desulfotignum* also constitutively expressed carbon monoxide dehydrogenase and acetyl-CoA synthase (*cooS/acsA*), the key genes in the acetyl-CoA pathway for acetate oxidation or carbon dioxide fixation (Fig. 3). Both, autotrophic growth and heterotrophic growth using genes of the acetyl-CoA pathway has been previously described for *D. balticum* (Kuever *et al*., 2001). Together, the high transcriptional activity of these genes (Fig. 3, Dataset 1) indicated that the organisms most likely fermented acetate to H2 and CO2, and then used the H2 for sulfate reduction (Kuever *et al*., 2001). However, it cannot be ruled out that the organisms grew chemolithoautotrophically, despite the excess of organic carbon sources, whichwould be counter-intuitive and merits further investigation. The *Desulfovibrio* population (bin F) also transcribed Ni/Fe hydrogenases (*hyb/hyd*) and appeared to consume hydrogen. It also transcribed genes for formate-hydrogen lyase (*hycE*) and formate oxidation (Fig. 3), consistent with the physiology of many *Desulfovibrio* species (Barton and Fauque, 2009). The coexistence of *Desulfotignum* and *Desulfovibrio* populations in each treatment of the experiment, revealed two stable ecological niches for sulfate reducers in our cultures.

#### Obligate fermenters

All *Clostridiales* (bin G-K, Fig. S7), *Spirochaetales* (bin L-M, Fig. S7), and *Anaerolineales* (bin P) were strictly fermentative, based on their gene content and transcriptional activity. They transcribed thioredoxins, peroxiredoxins and rubredoxins to protect their enzymes against oxidative stress during oxygen or nitrate treatments (Fig. 4B). The organisms transcribed hydrogen-producing hydrogenases and their associated electron transfer apparatus, but lacked a respiratory chain. All fermenters transcribed genes for electron transport complexes (*rnf*), which apparently enabled them to harness a proton/sodium motive force to reduce ferredoxins by oxidizing NADH. Glucose and amino acids supplied with the medium were the main substrates, as shown by highly transcribed sugar and amino acid transporters (Dataset 1). All *Firmicutes* (bin G – K) transcribed V-type and F-type ATP synthases. It was shown that F-type ATP synthases act as sodium pumps in certain *Clostridia* (Ferguson *et al*., 2006), so it is unclear whether these organisms harnessed a proton motive force to generate ATP. The three Spirochaetes only encoded a vacuolar type ATP synthase and are thus likely dependent on substrate level phosphorylation during fermentation. Transcription of acyl phosphatase and formate acetyltransferase (pyruvate-formate lyase) suggested that acetate and formate were end products of fermentation, in addition to hydrogen. All three end products seemed to be used by the two sulfate reducers, suggesting a syntrophic relationship between fermenters and sulfate reducers. The uncultured *Spirochaeta* bin L (Fig. S8) also transcribed genes to metabolize a large number of carbohydrates. The transcriptional activity indicated that this organism is able to import diverse sugars, into the cell and shuttle them into glycolysis or the pentose phosphate way (Fig. S10). Based on the transcription of key metabolic genes, the organisms affiliating with *Clostridiales* (bin G-I) seemed to have very similar physiologies, which was also the case for the organisms affiliating with *Spirochaeta* (bin L-N) (Fig. 3).

However, each population appeared to use slightly different glycosyl hydrolases (Table S1), and sets of genes involved in fermentation and energy conversion. For instance, the *Clostridiales* transcribed pyruvate synthase (*porC*), lactate dehydrogenase (*ldh*) and nitrite reductase (*nasD*), which the *Spirochaeta* did not transcribe. In turn, the Spirochaetes seemed to have a much higher expression of citrate synthase (*citA*) and isocitrate dehydrogenase (*icd*), key genes involved in the citric acid cycle (Fig. 3). The *Firmicutes* bin G exhibited high numbers of transcripts for amino acid importers and amino acid metabolism (e.g. glutamate dehydrogenase), whereas the *Firmicutes* bin K exhibited mainly transcripts of sugar importers and glycolysis. These differences in gene transcription may explain the observed coexistence of these organisms and hint towards metabolic complementation within the fermentative network.

#### Facultative aerobes and nitrate respirers

Populations affiliating with *Alphaproteobacteria* and *Gammaproteobacteria* (Fig. S9, bins A-C) were detected in the transient oxygen and nitrate cultures and were minor constituents in the sulfate-only cultures (Fig. 2). The transient exposure to oxygen and nitrate apparently selected for these organisms, which were capable of respiration. Genes encoding respiratory complexes I-IV and genes of the citric acid cycle were present and actively transcribed in the *Rhodobacterales* (bin A) and *Alteromonadales* (bin B). Compared to the fermenters, the respiratory organisms showed low transcriptional activity of sugar and amino acid transporters. Thus, it is likely that the respiratory organisms mainly used fermentation products, such as acetate, as electron donors. Hydrogen did not seem to be a major energy source for these organisms, as transcriptional activity of hydrogenases was not detected. In contrast, the *Rhodobacterales* actively transcribed all *sox* genes that are needed for sulfide and sulfur oxidation. Both organisms transcribed genes involved in polyhydroxybutyrate (PHB) and polyphosphate metabolisms. This indicates that PHB may have accumulated under anoxic conditions driven by polyphosphate hydrolysis, and was oxidized under oxic conditions, a well-known strategy for biological phosphorus removal (Wu *et al*., 2010). Indeed, in the *Rhodobacterales*, polyphosphate kinase and poly-beta-hydroxybutyrate polymerase were down-regulated during the period of air supply (Dataset 1).

The population related to *Desulfuromusa bakii* (bin E), did not have or transcribe *dsr* genes and was apparently not performing sulfate reduction. This organism were only selected in cultures with transient nitrate supply and showed a strong global transcriptional response to nitrate availability. In response to nitrate, it transcribed genes for citric acid cycle enzymes, complex I, nitrate-induced formate dehydrogenase (*fdn*), periplasmic nitrate reductase (*nap*), and pentaheme nitrite reductase (*nrf*). It likely performed nitrate ammonification with substrates such as amino acids, acetate and formate. *Desulfuromusa bakii* and related bacteria are known as sulfur-reducing, and often facultatively fermentative bacteria (Liesack and Finster, 1994). Hence, in the absence of nitrate the organisms selected here may also have performed fermentation of amino acids and/or dicarboxylates.

### Ecophysiology of *Candidatus* Sabulitectum silens

We also detected an organism (bin O) that affiliated with the candidate phylum *Fermentibacteria* (formerly candidate division Hyd24-12) (Kirkegaard *et al*., 2016). These organisms were present in all cultures, but were only abundant in the untreated cultures that were not exposed to oxygen or nitrate (Fig. 2). The contigs of this bin were very long (up to 538 kb; N50: 222 kb), the provisional genome had a size of 2.9 Mb and was inferred to be 77% complete (Table S2). Annotation of the genes encoded on the contigs of bin O suggested that the organisms have a typical gram-negative cell envelope with a complete peptidoglycan biosynthesis pathway and an active outer membrane transport system (*tonB*/*exbBD*). Glycolysis and the non-oxidative pentose phosphate pathway were complete (Fig. 5). The presence of largely complete operons coding for genes involved in lipid biosynthesis, cofactor biosynthesis, amino acid metabolism, and nucleotide metabolism indicated that these bacteria are likely not dependent on others for the generation of the major cellular building blocks. The organism transcribed an H^+^/Na^+^-translocating V-type ATP synthase as well as numerous protein complexes that translocate sodium ions across the cell membrane, such as an electron transport complex protein (*rnf*), a NADH-oxidoreductase (*ndh*), and a Na^+^-translocating decarboxylase (*oad/gcd*). This combination of proteins indicated that the organism was able to synthesize ATP using a sodium motive force (Mulkidjanian *et al*., 2008). However, the organism lacked a complete citric acid cycle and a respiratory chain. Single genes for flagellar biosynthesis and twitching motility were transcribed, yet the pathways for motility were incomplete (Fig. 5). Bin O lacked many of the mechanisms for oxidative and general stress protection (Fig 4B), which may explain its low abundance in the oxygen and nitrate treated cultures. The metagenome and metatranscriptome indicated that the organism is a non-motile, strictly anaerobic, obligate fermenter. We propose to name it *Candidatus* Sabulitectum silens (gen. *et* sp. nov.; *sabulum* (lat.) – sand; *tectus* (lat.) – covered, roofed; *silens* (lat.) – still, silent). In addition, we propose the new family *Ca*. Sabulitectaceae (fam. nov.) within the order *Ca*. Fermentibacterales (Fig. 6). The phylum *Fermentibacteria* belongs to the Fibrobacteres-Chlorobi-Bacteroidetes superphylum (Fig. S11). The *Fermentibacteria* comprise one class, one order, four families and at least nine genera (Fig. 6). The four families were previously indicated as four distinct clades (Kirkegaard *et al*., 2016).

**Figure 5:**
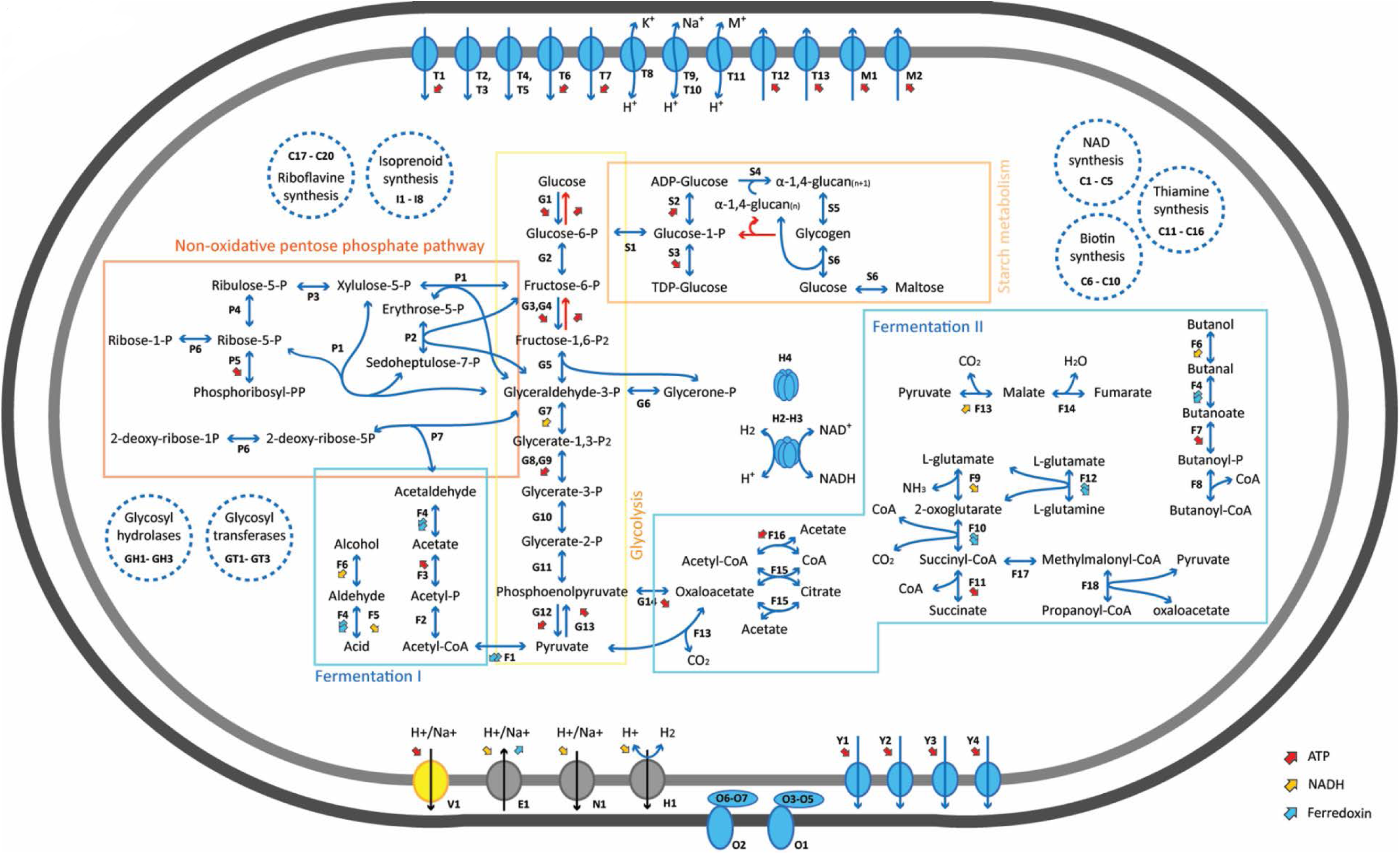
Metabolic map of *Ca*. Sabulitectum silens (bin O) showing central pathways that the organism transcribed in the sulfate-only treatment (Con-5, Con-6). Transcribed genes are shown as blue arrows, genes of annotated pathways that were not detected as red arrows. Enzymes are abbreviated with letters, the full list as well as further metabolic pathways are provided in Table S3. Dashed blue circles depict additional pathways that were detected.

**Figure 6:**
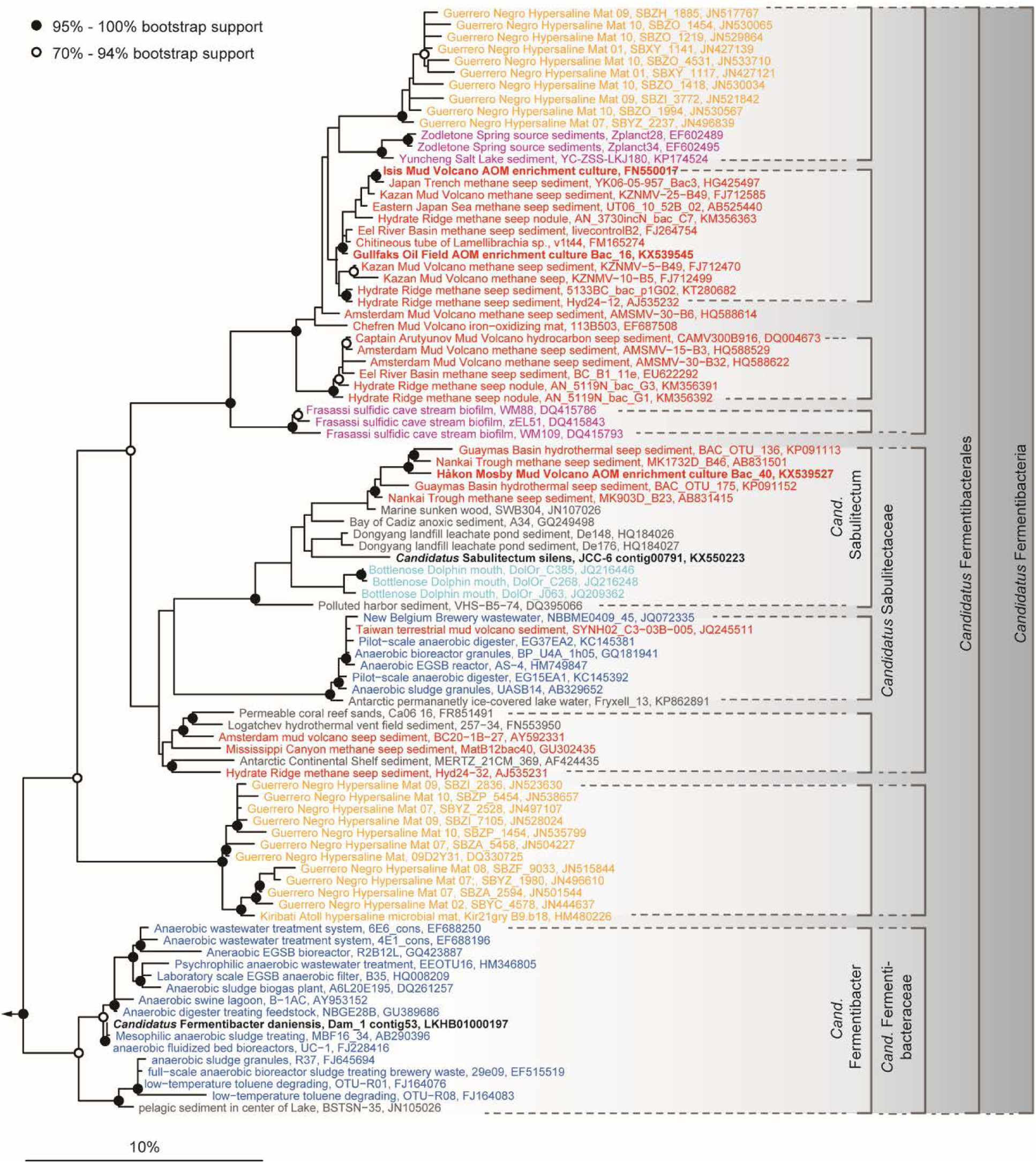
Phylogenetic tree of the phylum “*Ca*. Fermentibacteria”, showing the affiliation of all publicly available, non-redundant 16S rRNA gene sequences, including the provisional species *Ca*. Sabulitectum silens and *Ca*. Fermentibacter daniensis (black). The phylum comprises one class (at a threshold sequence identity of 78.5%), one order (at 82%), four families (at 86.5%) and at least nine genera (at 94.5%). The origin of the sequences is color-coded (red: methane seeps; red bold: anaerobic methanotrophic enrichment cultures; orange: hypersaline mats; pink: springs; light blue: dolphin, dark blue: anaerobic digesters: grey: other) and indicates niche-differentiation among *Fermentibacteria*. An extensive list of ecosystems harboring *Fermentibacteria* is provided in the Supplementary Results. Phylogeny is based on the SILVA small subunit ribosomal database SSURef 123.1 (released 03/2016). The scale bar shows estimated sequence divergence. *Fermentibacteria* sequence alignments and phylogeny are provided as an ARB database (Dataset 3). The parameters that were used to compile the sequence database are described in the Supporting Information.

The nearest relative of *Ca*. Sabulitectum silens is the recently described *Ca*. Fermentibacter daniensis, an anaerobic fermenter that is possibly involved in the sulfur-cycle (Kirkegaard *et al*., 2016). In contrast to *Ca*. Fermentibacter, *Ca*. Sabulitectum did not seem to possess or transcribe genes for sulfhydrogenases, despite the presence of sulfur in the cultures. Overall, both organisms appear to have similar lifestyles based on their transcriptional activity, despite their phylogenetic distance, suggesting that this lifestyle might be common among the phylum *Fermentibacteria*. Thus, it is not surprising that, so far, the phylum comprises sequences that almost exclusively originated from anoxic, organic and/or methane-rich ecosystems (Fig. 6), including sulfidic cave biofilms (Macalady *et al*., 2006), sulfur-rich springs (Elshahed *et al*., 2007), methane seeps (Ruff *et al*., 2015; McKay *et al*., 2016; Trembath-Reichert *et al*., 2016), mud volcanoes (Pachiadaki *et al*., 2011; Chang *et al*., 2012), methane hydrates (Mills *et al*., 2005), marine sediments (Schauer *et al*., 2011), coral reef sands (Schöttner *et al*., 2011), microbial mats (Harris *et al*., 2013; Schneider *et al*., 2013), marine sponges (Simister *et al*., 2012) and anaerobic digesters (Nelson *et al*., 2012; Kirkegaard *et al*., 2016). The physiology that *Ca*. Sabulitectum exhibited in our cultures (Fig. 3, 4B, 5) suggests that *Fermentibacteria* are strict anaerobes that produce hydrogen and acetate from the fermentation of amino acids and sugars, in these ecosystems.

## Conclusion

The transient exposure to oxygen or nitrate changed the microbial community structure, and impacted the magnitude of net sulfide production as a community function, yet had a minor effect on microbial community composition. This shows that the communities of Janssand intertidal sediments contained organisms that were well adjusted for each of these scenarios, diverting the flow of carbon and energy through the trophic network based on the available electron acceptors. The treatment with oxygen or nitrate did not cause the community to shift to an alternative stable state (Shade *et al*., 2012). Community stability during the exposure to oxygen or nitrate was enabled by the increased expression of genes involved in oxidative and general stress protection. The stable coexistence of several fermenters and sulfate reducers with nitrate reducers or aerobic respirers supports the recent finding that microbial communities are assembled based on rules that go beyond those of the classical redox tower (Chen et al., 2017).

## Materials and Methods

### Sampling site and inoculum for enrichment experiments

Sediment was sampled from the upper part of the intertidal back-barrier flat Janssand, in the German Wadden Sea (53.73515 N, 07.69913 E) in June 2012. The top 2 cm of sediment was collected with a flat trowel during low tide. After transport of the sediment to the laboratory, an equal volume of sterile artificial seawater (Red Sea Salt, 33.4 g l^-1^; http://www.redseafish.com) was added to the sediment and stirred vigorously. The sediment was allowed to settle briefly, and the liquid was transferred into (1 l) glass bottles that were closed with rubber stoppers and of which the headspace was exchanged with argon. The liquid was kept at 4°C for 2 days and then used as inoculum.

### Continuous culture setup and medium

Six continuous cultures were set up and maintained for 350 days. Each glass vessel (DURAN, GLS 80, 500 ml) was filled with 0.4 l inoculum, fitted with tubes for in- and outflowing medium as well as in- and outflowing gas, and was stirred at 200 to 400 rpm. The medium supply rate was 0.17 l day^-1^, resulting in a dilution rate of 0.36 – 0.4 day^-1^. The anoxic medium consisted of Red Sea Salt artificial seawater (33.4 g l^-1^), containing 28 mM sulfate, supplemented with 20 C-mM organic carbon (1.1 mM D-glucose, 1.7 mM acetic acid, and 0.4 mM amino acids), 0.2 mM phosphate and trace elements (for details see Supporting Information). The culture pH was measured off line (Mettler Toledo, Five Easy ^TM^) and was in the range of pH 7.5 to 7.8. The OD_600_ of all cultures was monitored off-line spectrophotometrically (Thermo Scientific Genesys 10S UV-Vis). Sulfide concentration in the culture was measured using the Cline method (1969). After 21 days, the headspace of two of the cultures (Oxy-1 and Oxy-2) was oxygenated twice daily, by supplying air (1 l min^-1^) for 5 min. The air was removed after 30 min by supplying Argon (1 l min^-1^) for 5 min. This procedure was repeated every 12 h for the remainder of the experiment. In parallel, nitrate was supplied twice daily to two cultures (Nit-1 and Nit-2) by supplying a nitrate solution (1.4 ml min^-1^, 20 mM NaNO_3_ dissolved in artificial seawater) for 7 min, every 12 hours. Two other cultures (Con-1 and Con-2) only received sulfate as electron acceptor. During aeration, the oxygen concentration in the culture liquid was measured with Optical Oxygen Meter – FireSting O2. At the same time, we measured off-line the hydrogen sulfide (Cline, 1969) and sulfur concentrations (Kamyshny Jr and Ferdelman, 2010) in the cultures. In addition, after 327 days the nitrate in the medium was replaced with ^15^ N-nitrate by direct injection of 10 ml of 20 mM ^15^ N-nitrate and the production of ^15^ N-nitrogen gas was measured off-line by mass spectrometry (GAM 400, InProcess Instruments, Bremen, Germany) using 0.5 ml headspace samples. Nitrate in the culture liquid was determined as previously described (Hanke *et al*., 2014).

### Metagenomics

On day 311 (Oxy-1 and Oxy-2), day 327 (Nit-1 and Nit-2), and day 300 (Con-1 and Con-2) of the experiment, we extracted nucleic acids from 10 ml samples of all six cultures as previously described (Zhou *et al*., 1996), after incubation with lysozyme (2.5 mg ml^-1^) and RNAse (0.1 mg ml^-1^). For metagenome shotgun sequencing, 1.5 μg of the extracted DNA was mechanically fragmented using Nebulizers (Roche; 32 psi; 3 min, 500 μl nebulization buffer). The fragmented DNA was purified using MinElute PCR purification columns (Qiagen) and eluted in 50 μl Tris-EDTA buffer (Life Technologies). The entire eluate was used for the preparation of barcoded Personal Genome Machine (PGM) sequencing libraries with the Ion Xpress^TM^ Plus gDNA Fragment Library Preparation kit (Life Technologies). Library insert sizes were between 350 and 400 base pairs (bp). The libraries were sequenced with the PGM on a 318 Chip, using the chemistry for 400 bp libraries. Base calling was performed with the Torrent Suite software v3.6 or v4.0.2, with default settings. Sequence reads were assembled with the Newbler assembler v2.8 with default settings for genomic DNA assembly for non-paired reads. Assembled DNA sequences were binned based on multivariate statistics of tetranucleotide frequencies with MetaWatt v2.1 (Strous *et al*., 2012) (Fig. S1, Table 1). Phylogenetic profiles of the bins were obtained by analyzing all open reading frames encoded on the contigs using blast and a database that contained a representative of every genus with a publicly available, complete or draft whole genome sequence. Genome completeness and contamination were evaluated by detection of a set of 139 conserved single copy genes (Campbell *et al*., 2013) with Hidden Markov Models (HMMER 3.1) and by detection of transfer RNA genes (Laslett and Canback, 2004). Percentage completeness was calculated as the number of conserved single copy genes (CSCG) detected, divided by the total number of CSCG. Percentage contamination was calculated as the number of CSCG present in >1 copy, divided by the number of CSCG detected. Due to frameshift errors resulting from Ion Torrent sequencing it was not possible to use CheckM (Parks *et al*., 2015) to estimate genome completeness and contamination. Note that, using the above described method, the completeness values reported likely underestimate the actual completeness of the bins. Genes present in each bin were annotated with Prokka v1.9 (Seemann, 2014). Each bin constitutes a provisional whole genome sequence of a microbial population (Fig. S1). Bins denoted by the same letter across all cultures (e.g. bin D) represent a genetically identical population, or nearly identical populations, since all six continuous cultures were inoculated with the same sediment sample. Bin abundances over all samples were estimated based on coverage and bin size, by mapping the sequence reads to the contigs that made up each associated bin. The abundance of the enriched organisms in the inoculum was estimated by mapping the reads of four Janssand sediment metagenomes (Sequence Read Archive accession numbers SRR577219, SRR577220, SRR577221, SRR577224) to the bins using BBMap (github.com/BioInfoTools/BBMap) and the parameters “maxlen 500, minid 0.98”.

### Metatranscriptomics

Parallel to DNA extraction, RNA was extracted from a 2 ml sample of all six cultures (for details see Supporting Information) on day 311 (oxygen treated cultures), on day 327 (nitrate treated cultures) and on day 300 (untreated control cultures). For those cultures with cyclic oxygen or nitrate supply, RNA was extracted an hour before the treatment and immediately after the treatment subsided, i.e. when oxygen and nitrate concentrations had decreased to background values, 30 min and 240 min after the treatment commenced, respectively (Fig. S3 C, D). Ribosomal RNA was depleted from purified RNA (3–5µg) using the Ribo-Zero rRNA removal kit (Bacteria, Epicentre, Madison,WI, USA). Libraries were prepared with the Ion Total RNA-Seq Kit v2 (Life Technologies) following the protocol for whole transcriptome library preparation. Transcriptional activities for each gene (Dataset 1) were determined by Ion Torrent sequencing of cDNA obtained from extracted RNA and subsequent mapping of the cDNA reads to the annotated contigs with BBMap v32 (http://sourceforge.net/projects/bbmap/). Reported activities were calculated by dividing the number of mapped reads/gene length by the total number of reads mapped to coding sequences of the bin/total length of all coding sequences of the bin.

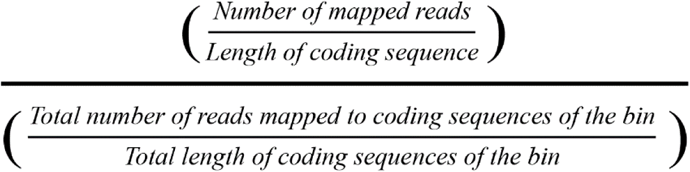

This way, the average transcriptional activity equals 1.0 and hence the bins can be compared within and between the treatments. 16S rRNA gene sequences were detected with Hidden Markov Models (www.github.com/Victorian-Bioinformatics-Consortium/barrnap) and, independently, reconstructed with Emirge (Miller *et al*., 2011). 16S rRNA gene sequences were linked to bins as previously described (Kraft *et al*., 2014).

### Phylogenetic tree reconstruction

The 16S rRNA gene based phylogenetic trees were generated using near full-length sequences (>1300 bases) of the non-redundant SILVA small subunit reference database (release 123.1; March 2016) (Quast *et al*., 2013) and the software ARB (Ludwig *et al*., 2004). Sequences were aligned using SINA (Pruesse *et al*., 2012) and the alignment was manually optimized according to the rRNA secondary structure, resulting in high-quality alignments of 1267–1287 bases length. We used a maximum-likelihood algorithm (PHYML) with a positional variability filter, excluding highly variable regions, and 100 bootstrap iterations. Phylogenetic levels were calculated based on phylogenetic distance using the clustering tool as implemented in ARB. Threshold sequence identity for genus (94.5%), family (86.5%), order (82.0%), class (78.5%) and phylum (75.0%) were chosen according to the latest taxonomic threshold recommendations (Yarza *et al*., 2014).

### Sequence data accession

16S rRNA gene sequences are archived under the accession numbers KX550146 – KX550265 (Janssand continuous cultures) and KX539512-KX539546 (Seep sediment enrichments). Metagenomic and metatranscriptomic sequencing raw data, as well as the assembled contigs and the *Ca*. Sabulitectum silens draft genome, are archived under the SRA Bioproject PRJNA305678 and the BioSamples SAMN04331582-SAMN04331591

## Acknowledgements

We gratefully acknowledge Brandon Kwee Boon Seah for support with data processing, Miriam Sadowski, Veronika Will, and Gunter Wegener for assistance with AOM enrichment cultures, and Kirsten Imhoff for sulfide measurements. We thank Fridjof Boness for help with nomenclature as well as Katrin Knittel, Emmo Hamann and Manuel Kleiner for discussions. ZB was supported by a Yousef Jameel Scholarship. SER was supported by an AITF/Eyes High Postdoctoral Fellowship. The research was funded by an ERC starting grant (MASEM, 242635), a Campus Alberta Innovation Chair and a NSERC Discovery grant awarded to MS, the German Federal State North Rhine Westphalia and the Max Planck Society.

